# Full-length mRNA sequencing uncovers a widespread coupling between transcription and mRNA processing

**DOI:** 10.1101/165092

**Authors:** Seyed Yahya Anvar, Guy Allard, Elizabeth Tseng, Gloria Sheynkman, Eleonora de Klerk, Martijn Vermaat, Raymund H. Yin, Hans E. Johansson, Yavuz Ariyurek, Johan T. den Dunnen, Stephen W. Turner, Peter A.C. 't Hoen

## Abstract

The multifaceted control of gene expression requires tight coordination of regulatory mechanisms at transcriptional and post-transcriptional level. Here, we studied the interdependence of transcription, splicing and polyadenylation events on single mRNA molecules by full-length mRNA sequencing. In MCF-7 breast cancer cells, we found 2,700 genes with interdependent alternative transcription, splicing and polyadenylation events, both in proximal and distant parts of mRNA molecules. The analysis of three human primary tissues revealed similar patterns of interdependency between transcription and mRNA processing events. We predict thousands of novel Open Reading Frames from the sequence of full-length mRNAs and obtained evidence for their translation by shotgun proteomics. The mapping database rescued 358 previously unassigned peptides and improved the assignment of others. By recognizing sample-specific amino-acid changes and novel splicing patterns, full-length mRNA sequencing improved proteogenomics analysis of MCF-7 cells. Our findings demonstrate that our understanding of transcriptome complexity is far from complete and provides a basis to reveal largely unresolved mechanisms that coordinate transcription and mRNA processing.

## INTRODUCTION

The formation of a mature messenger RNA (mRNA) is a multi-step process. In higher eukaryotes, variations in each of these steps, including alternative transcription initiation, differential splicing of exons, and alternative polyadenylation site usage, change the content of the mature transcript. The multitude of transcripts arising from these events offers an enormous diversity of protein isoforms that can be produced from a single gene locus. Tight regulation and coordination of these processes ensures the production of a (limited) set of cell-, tissue-and condition-specific transcript variants to meet variable cellular protein requirements^1-4^. Whether or not these processes are co-transcriptionally linked is currently largely unknown, as are the mechanisms that couple transcription with 5’ end capping, splicing, and 3’ end formation (reviewed in 5). Thus, resolving full transcript structures and accurate quantification of the abundance of alternative transcripts are important steps towards the detection and understanding of these mechanisms.

RNA sequencing (RNA-Seq) has become a central technology for deciphering the global RNA expression patterns. However, reconstruction and expression level estimation of alternative transcripts using standard RNA-Seq experiments is limited and prone to error due to relatively short read length (typically up to 150 nucleotides) and required amplification steps of second-generation sequencing technologies^6^,^7^. It is apparent that single-molecule long reads that capture the entire RNA molecule can offer a better understanding of the rich patterns of alternative transcription and mRNA processing events and, hence, the underlying biology.

Despite a number of studies that have pursued long read sequencing to connect different exons or even capture entire transcripts with a rather limited sequencing depth^6^,^8-14^, the coupling between transcription and mRNA processing has not been extensively studied. Here, we investigate the global pattern of coupling between transcription, splicing and polyadenylation in MCF-7 human breast cancer cell line and three human tissues, which are deeply sequenced using the single-molecule real-time Pacific Biosciences RSII sequencing platform. We show that transcription and mRNA processing are tightly coupled and that such interdependencies can be found across the entire RNA molecule and across large intra-molecular distances. We demonstrate that transcript identification and understanding of coupling between processes that are involved in the formation of these transcripts is far from complete, even in well-characterized human cell lines such as MCF-7. This study provides an in-depth view of the true complexity of the transcriptome and, for the first time, shows the tight and global interdependency between alternative transcription, splicing and polyadenylation. We also show the value of this resource in relation to translation and sample-specific survey of the proteome.

## RESULTS

### Detection of transcript variants and the associated interdependencies between alternative exons

To investigate the genome-wide coupling of transcription and mRNA processing events, full-length mRNAs from MCF-7 human breast cancer cells were sequenced on 147 SMRT cells using Pacific Biosciences RSII platform (**Supplementary Table 1**). Prior to sequencing, parts of the sequencing library were size selected to allow for a good representation of longer transcripts.

**Table 1.** Enrichment of MBNL binding site motifs in sequences upstream of alternative PAS with unknown poly(A) signal that are coupled with alternative TSS or alternative exons.

| Motifs | Source | Total | Random Set | P-value * | Coupled PAS | Not Coupled PAS | P-value § |
| --- | --- | --- | --- | --- | --- | --- | --- |
| AKCCTGG | DREME | 1,271 | 35 | 0 | 881 | 390 | 9.8E-44 |
| CTSCYB | Masuda, 2012 <sup>24</sup> | 898 | 708 | 2.6E-07 | 442 | 456 | 9.7E-01 |
| YGCY | Purcell, 2012 <sup>25</sup> | 2,961 | 3,139 | 1.0E-00 | 1,578 | 1,383 | 3.2E-02 |
| RSCWTGSK | Batra, 2014 <sup>23</sup> – MBNL1 | 145 | 93 | 4.1E-04 | 80 | 65 | 2.5E-01 |
| TGCYTSYY | Batra, 2014 <sup>23</sup> – MBNL2 | 95 | 55 | 6.5E-04 | 50 | 45 | 4.9E-01 |
| CWGCMWKS | Batra, 2014 <sup>23</sup> – MBNL3 | 1,306 | 139 | 4.7E-262 | 870 | 436 | 1.5E-32 |
| **Total PASs** |  | 6,979 | 6,979 | – | 3,614 | 3,338 | – |

\* The enrichment of binding motifs in sequences upstream of PASs without a known poly(A) signal were calculated by Fisher's exact test (one-sided). A randomly generated set was used as a background for enrichment analysis.
§ PASs without significant coupling were used as the background set to identify a binding site that is enriched in the coupled PASs without a known poly(A) signal.

Transcript structures were defined by applying the isoform-level clustering algorithm (ICE) on full-length reads, capturing the entire mRNA molecule (containing both 5’ and 3’ primer sequences). Transcript sequences were further polished using both full-length and partial reads (**Fig. 1A**). The analysis pipeline precisely determined the position of polyadenylation sites (presence of poly(A) tail in the sequence) and intron-exon boundaries, as evident from the presence of the canonical GU motif in 93% of donor splice sites and the canonical AG motif in 95% of acceptor splice sites. In fact, 90% of introns are defined by canonical splice-site motifs (GU-AG). In addition to 7,364 single-exon transcripts, the MCF-7 transcriptome consists of 11,350 multi-exon genes of which 69% produced multiple transcript structures (**Supplementary Fig. 1**). Transcripts range from 54bp to 10,792bp in length with an average of 82 supporting reads (**Supplementary Fig. 1**). Interestingly, 49% of identified transcripts in MCF-7 are identified as potentially novel in comparison with the GENCODE annotation (**Supplementary Table 2**).

**Table 2.**
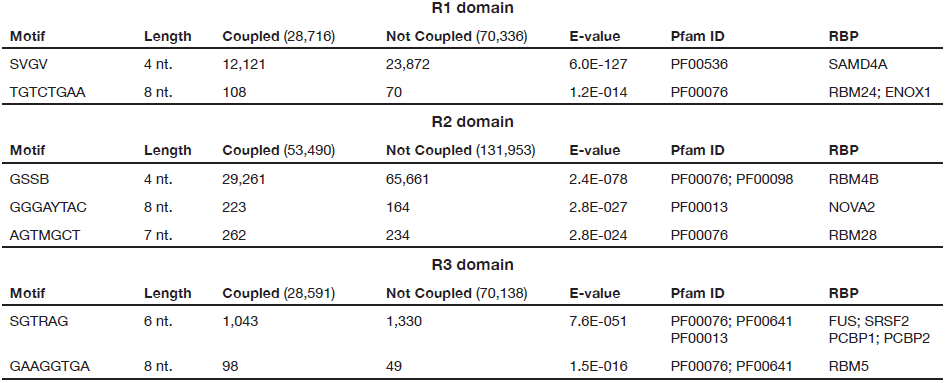
The RNA-binding protein motifs associated with alternative exons that are coupled to TSS, other alternative exons, or PAS.

| R1 domain |  |  |  |  |  |  |
| --- | --- | --- | --- | --- | --- | --- |
| Motif | Length | Coupled (28,716) | Not Coupled (70,336) | E-value | Pfam ID | RBP |
| SVGV | 4 nt. | 12,121 | 23,872 | 6.0E-127 | PF00536 | SAMD4A |
| TGTCTGAA | 8 nt. | 108 | 70 | 1.2E-014 | PF00076 | RBM24; ENOX1 |
| R2 domain |  |  |  |  |  |  |
| Motif | Length | Coupled (53,490) | Not Coupled (131,953) | E-value | Pfam ID | RBP |
| GSSB | 4 nt. | 29,261 | 65,661 | 2.4E-078 | PF00076; PF00098 | RBM4B |
| GGGAYTAC | 8 nt. | 223 | 164 | 2.8E-027 | PF00013 | NOVA2 |
| AGTMGCT | 7 nt. | 262 | 234 | 2.8E-024 | PF00076 | RBM28 |
| R3 domain |  |  |  |  |  |  |
| Motif | Length | Coupled (28,591) | Not Coupled (70,138) | E-value | Pfam ID | RBP |
| SGTRAG | 6 nt. | 1,043 | 1,330 | 7.6E-051 | PF00076; PF00641<br>PF00013 | FUS; SRSF2<br>PCBP1; PCBP2 |
| GAAGGTGA | 8 nt. | 98 | 49 | 1.5E-016 | PF00076; PF00641 | RBM5 |

**Figure 1.**
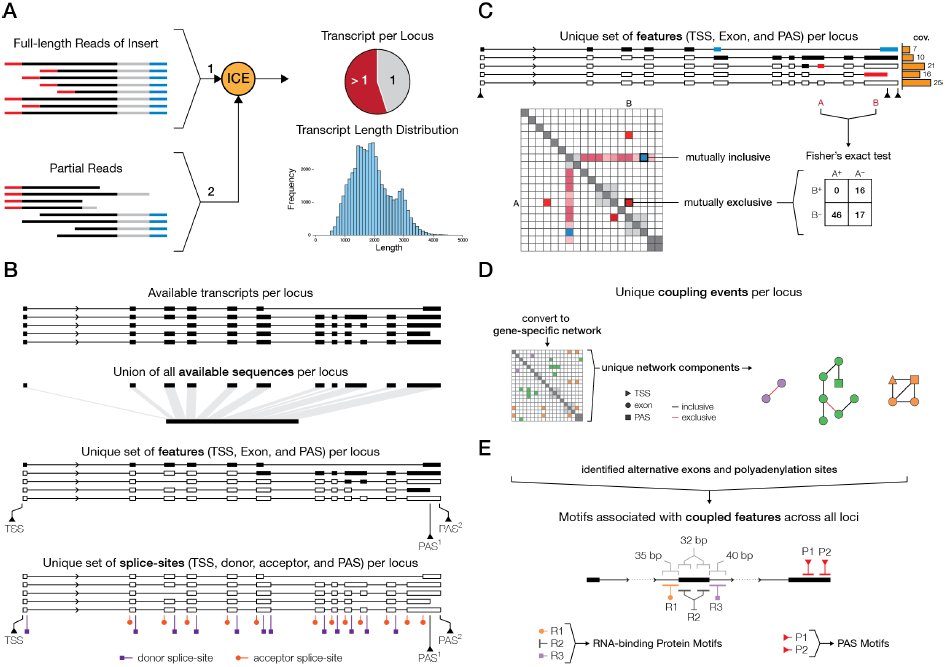
Schematic overview of the approach to characterize the interdependencies between mRNA transcription and processing events. **A)** Identified full-length reads (reads with RNA inserts between 5’ and 3’ primers) are clustered into unique transcript struc-tures using the ICE algorithm and further polished using the partial reads, where one of the primer sequences is missing. **B)** Based on available transcripts per locus, available sequence (union of all exonic sequences that are observed at each locus) and unique set of features and splice-sites are identified. Features sets comprise of unique transcription start sites (TSS), exons, and polyadenylation sites (PAS). The unique set of splice sites consists of unique donor and acceptor splice-sites as well as all alternative TSSs and PASs. **C)** The survey of coupling events is done by performing all possible pairwise tests between unique features in genes. The sum of the coverage of all transcripts that support the inclusion or exclusion of each pair is used in a contingency table to perform a Fisher’ s exact test for statistical significance. The odds ratio (OR) is used to differentiate between mutually inclusive and exclusive coupling. **D)** Set of interdependent coupling events were identified based on networks of coupling between features in each gene. Nodes represent features and links depict the mutual inclusivity (black edges) or mutual exclusivity (red edges) of each feature pair. Unique network components can thereby be filtered based on the type of interaction: mutual inclusive or mutual exclusive coupling events. **E)** For all alternative exons that show significant linkage, a motif search is performed to assess the enrichment of specific RNA-binding protein motifs. For all alternative exons, 35bp intronic sequences upstream of the acceptor site are defined as R1 domain (depicted in orange), 32bp exonic sequences downstream of the acceptor site and upstream of the donor site are defined as R2 domain (depicted in dark grey), and 40bp intronic sequences downstream of the donor site are defined as R3 domain (depicted in purple). 35bp sequence upstream of each PAS (depicted in red) is searched for the presence of canonical and non-canonical poly(A) signals.

The gene expression levels measured in full-length mRNA sequencing data showed significant Spearman correlations (correlation coefficient greater than 0.74) with 5 publicly available RNA-Seq datasets that were generated on Illumina HiSeq2000 or HiSeq2500 platforms (**Supplementary Fig. 2A**). Differences in library preparation protocols, presence of fewer duplicates and uniformity of coverage in the PacBio data as well as contrast in sequencing dynamics contribute to minor differences observed in estimated gene expression levels^15^ Although we observed some inter-dataset differences in detected genes, for most genes, the results from all datasets were in concordance (**Supplementary Fig. 2B**). Transcript lengths and loading bias did not significantly contribute to inter-platform differences for gene detection as similar results were also found for intra-platform comparisons (**Supplementary Fig. 2C**). Thus, the full-length mRNA sequencing data can be reliably used for locus-specific quantification of transcript abundance in MCF-7 cells.

**Figure 2.**
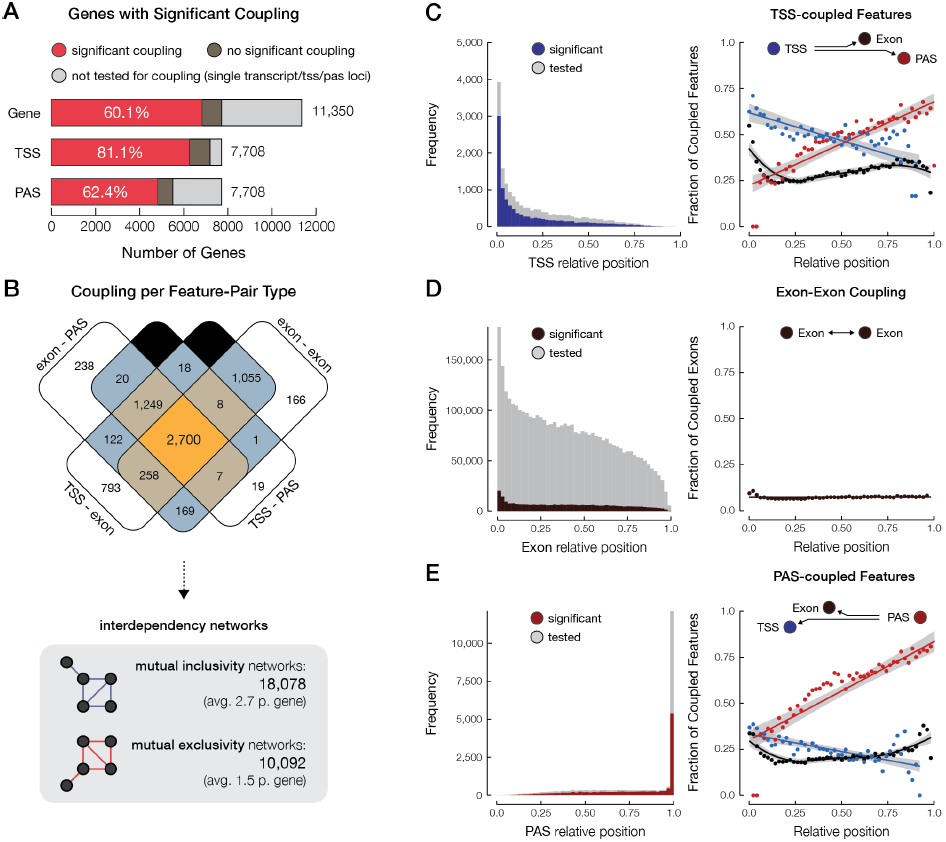
Alternative transcription, splicing, and polyadenylation are highly interdependent. **A)** Bar charts illustrate the number and proportion of genes that show significant coupling. Genes with TSS- or PAS-coupled features are also presented. **B)** Venn diagram shows the number of genes with various types of coupling representing interdependencies between different alternative processes. The total number of mutually inclusive and exclusive networks are also listed. **C)** Histogram of the relative positions of transcription start sites (TSS) with (blue) and without (grey) significant coupling to mRNA processing events. Relative positions are calculated based on the length of the total exonic sequence at each locus. Scatter plot shows the fraction of significantly coupled TSSs (blue) to alternative exons (black) and PASs (red), plotted at each relative position.**D)** Histogram of the relative positions of alternative exons with (brown) and without (grey) significant coupling to other exons. Scatter plot shows the fraction of significantly coupled exons to other exons, plotted at each relative position. **E)** Histogram of the relative positions of polyadenylation sites (PAS) with (red) and without (grey) significant coupling to alternative transcription and splicing events. Scatter plot shows the fraction of significantly coupled PASs (red) to alternative TSSs (blue) and exons (black), plotted at each relative position. For plots depicting the percentage of linked features per position, the bin size of 0.02 was used.

To detect and characterize the dependency between transcription and mRNA processing events, we designed the following analysis strategy (**Fig. 1**): For each gene, the union of all exonic sequences was considered as the available sequence and the union of all unique transcription start sites (TSSs), exons (defined as having distinct donor and acceptor splice sites), and polyadenylation sites (PASs) was used as a set of available features (**Fig. 1B**). Mutual inclusivity or exclusivity of all possible combinations of features was assessed based on the number of reads that support the inclusion or exclusion of each pair of features. Subsequently, we applied a Fisher’s exact test to evaluate statistical significance of the interdependency between a pair of features (**Fig. 1C**; also see Methods). It is important to note that as features may be coupled to a few other features, the actual coupling events can be summarized into a series of network components within a gene-specific interaction network to capture the independent coupling events (**Fig. 1D**). These components can be summarized based on the level of connectivity or mutual inclusivity or exclusivity within each network to construct subnetworks. We subsequently searched the sequences containing the coupled alternative exons or poly(A) sites for enriched sequence motifs and tested whether they contain motifs of known RNA-binding proteins (**Fig. 1E**).

### General properties of coupling in human MCF-7 transcriptome

The MCF-7 transcriptome data consist of 11,350 multi-exon genes that present 3,532,796 combinations of features (TSSs, exons, and PASs). Most combinations represent exon-exon pairs as many loci contain only a single TSS or PAS whereas all loci are multi-exonic (**Supplementary Fig. 3**). Since the test is only valid for genes with multiple transcripts, only 7,708 genes and 3,055,099 pairs of features (TSSs, exons, and PASs) were included in the statistical analysis. Importantly, pairs of features that are naturally interdependent (such as overlapping features or PASs and the terminal exons) have been removed from the analysis.

**Figure 3.**
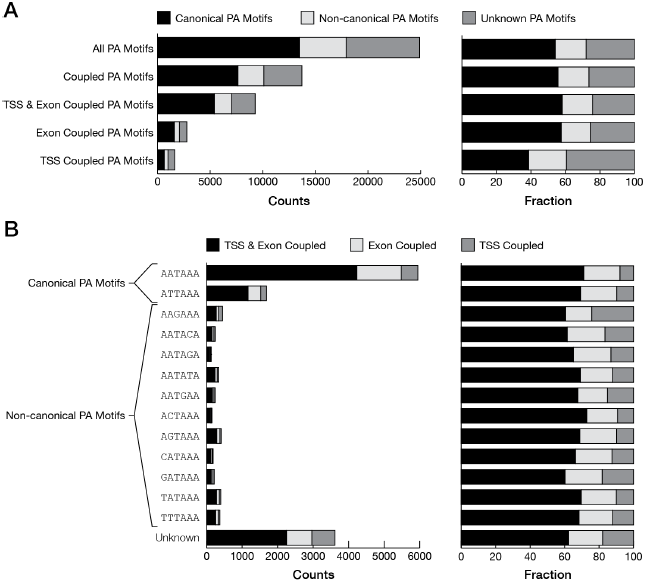
Alternative transcription start sites and exons are significantly associated with known and novel poly(A) signals. **A)** Bar charts show the number and relative proportion of PASs that are associated with canonical or non-canonical poly(A) signals for all PASs, PAS with significant coupling: alternative exon and/or TSS linked PASs.**B)** Bar charts represent the number and relative proportion of known and unknown poly(A) signals for TSS-linked, exon-linked, or TSS and exon linked PASs.

Almost ten percent of all feature pairs were significantly coupled (p-value < 1.4e-08, after Bonferroni correction for multiple testing). Generally, we observed large effect sizes for coupled features with almost equal distribution over mutually inclusive (52%) and mutual exclusive pairwise interdependencies (**Supplementary Fig. 4A**). Remarkably, we observed coupling between mRNA features in over 60% of all multi-exon genes (6,825 out of 11,350; **Fig. 2A**), represented by 18,078 mutually inclusive and 10,092 mutually exclusive subnetwork components (**Fig. 2B**). Particularly, alternative TSSs appear to have a significant impact on mRNA processing as over 80% of multi-transcript genes exhibit interdependency between the choice of TSS and alternative splicing. We found a substantial amount of interdependencies between all types of features (**Fig. 2B**; **Supplementary Fig. 4A**). Of the 6,825 genes with at least one coupling event, 2,700 (37%) showed interdependencies between all classes of features: alternative TSS linked to alternative exons, alternative exon to alternative exon linkage, alternative PAS linked to alternative exons, and alternative TSSs to alternative PASs. Thus, the deep sequencing of full-length mRNAs provided a first image of the large degree of coordination in the usage of alternative TSSs, exons and PASs, restricting the number of produced transcripts given the substantial amount of combinatorial possibilities.

**Figure 4.**
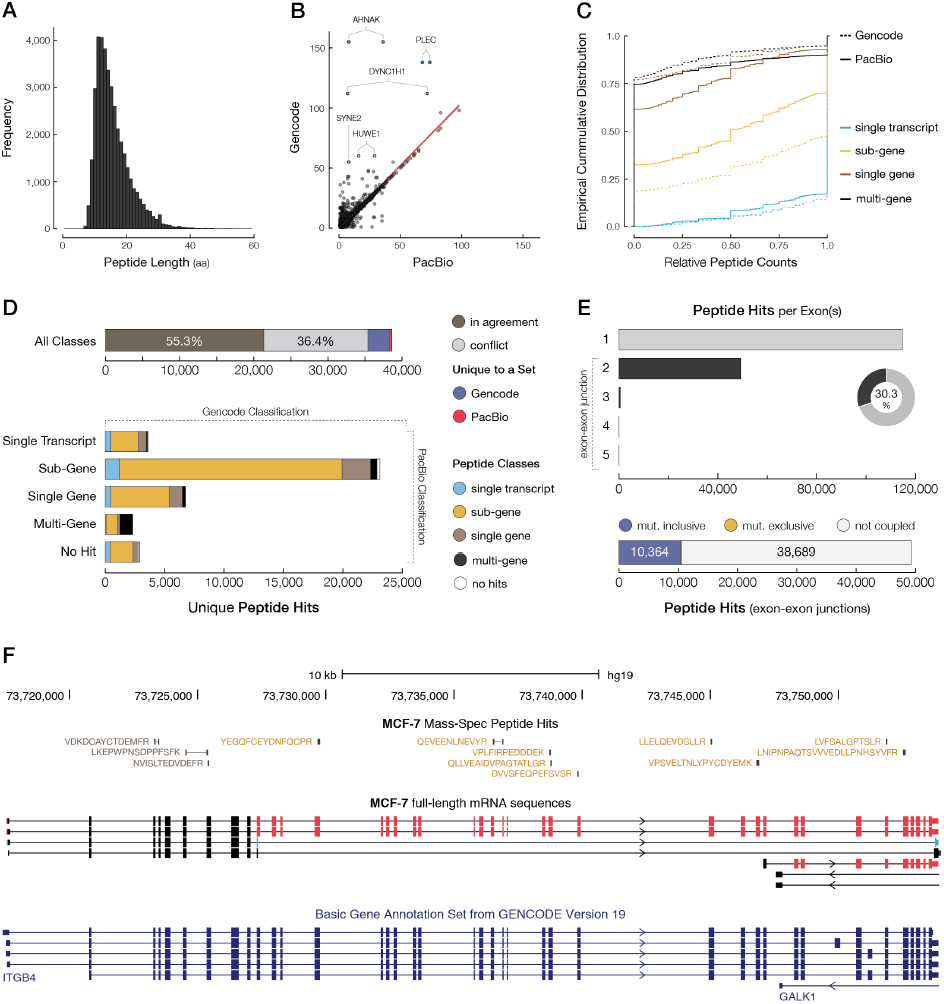
Comprehensive map of protein peptides supports novel alternative splicing events in full-length MCF-7 transcriptome. **A)** Histogram shows the distribution of peptide amino acid (aa) lengths that could be associated with either Gencode or PacBio transcript variants. **B)** Scatter plot illustrates the number of unique peptide hits per gene based on PacBio (x-axis) or Gencode annotation (y-axis). Each dot represents a single gene locus. **C)** Empirical cumulative distribution of relative peptide counts per gene for each peptide hit category. Genes with a single transcript annotation (single-transcript category) are shown in light blue. Multi-transcript genes with peptides matching to a subset of transcripts (sub-gene category) are shown in yellow. Multi-transcript genes with peptides matching to all annotated transcripts (single-gene category) are shown in brown. Multi-gene hits are shown in black. Dotted lines represent the cumulative distributions based on the Gencode annotation. **D)** Bar charts illustrate the comparison of Gencode- or PacBio-based classification of Peptides. **E)** Bar charts show the number of peptides derived from exon-exon junctions of transcripts. The number of peptides that match exon-exon junction of mutually inclusive (blue) or exclusive (yellow) exons.**F)** Peptides with different classification matching to multiple transcripts of ITGB4. Black peptides are single-gene hits whereas, based on full-length MCF-7 transcriptome data, yellow peptides are only associated with a subset of transcripts. Exons are colored based on coupling networks, shown in red and blue.

The length of individual transcripts was not associated with the likelihood of a significant coupling event in that transcript (**Supplementary Fig. 4B**). However, after examining the length of the union of exonic sequences per gene and the likelihood of observing coupling, we found that significant coupling events were enriched in genes with larger available exonic sequences (**Supplementary Fig. 4B**). Expectedly, larger exonic regions in each locus gives rise to a larger repertoire of possible transcripts, requiring more extensive regulation of the synthesis for transcripts containing different subset of features.

We also examined the effect of the relative position in the gene and the distance between features on the observed degree of coupling. As expected, most TSSs were located at the most 5’-end of genes. However, interdependency of alternative TSSs was observed across the entire gene (**Fig. 2C**, left panel). Alternative TSSs were preferentially coupled to alternative splicing events in relatively close proximity to the TSSs, near the 5’-end, as well as distal exons at the 3’-end (**Fig. 2C**, right panel). Nevertheless, examples of the coupling of alternative TSS and alternative exon usage across large distances, and spanning multiple exons were also frequently observed (**Supplementary Fig. 5,6**; *ALDOA* and *C1QTNF6*). More evidence for interactions across the entire length of genes comes from the significant coupling between TSS and PAS, the two most distant features in an mRNA molecule (**Supplementary Fig. 7**; *NCAPD2*).

The fraction of alternative exons was higher at the 5’-end; however, dependencies between multiple alternative splicing events were uniformly observed across the entire gene (**Fig. 2D**; **Supplementary Fig. 8**, *LMNA*). Despite the uniform distribution of exon-exon coupling events and the presence of distant coupling events (**Supplementary Fig. 9**, *RELA*), most interdependent alternative splicing events were between nearby or neighboring exons (**Supplementary Fig. 10**, *CALU*).

Like alternative transcription start sites, coupling events linked to alternative PAS usage were found across the entire gene (**Fig. 2E**). In concordance with published literature^16-18^, alternative PAS usage was preferentially coupled to nearby alternative exons (**Fig. 2E**). Nevertheless, a substantial proportion of PASs was coupled to alternative exons in more 5’-regions of genes.

We performed Sanger sequencing to independently verify identified coupling events for a set of gene loci. Due to limited range of alternative technologies such as Sanger sequencing, only relatively close coupling events could be assessed. In most cases, the Sanger sequencing results were in full concordance with the coupling events identified using full-length mRNA sequencing (**Supplementary Fig. 5,6,8,9**). Next, we carried out a single-molecule RNA in situ fluorescence (smRNA FISH) co-localization approach^19^,^20^ to examine the alternative splicing events that were identified by full-length RNA sequencing and confirmed by Sanger sequencing. Four probe sets were designed to detect different segments of the *CALU* mRNAs at single-molecule level. The full CALU_E probe set covers the common exons on the full-length variants, whereas the 9-oligo CALU_E4 and CALU_E5 sets were designed to specifically hybridize to either exon 4 or 5. The signal from the common exon probes was easily detected (**Supplementary Fig. 10**), and could be used for co-localized signals from the exon 4, exon 5 and intron 1 sets. The average numbers and distribution of signals revealed that cytoplasmic mRNAs predominantly either include exon 4 (~50 copies per cell) or exon 5 (~10 copies per cell), followed by mRNAs that do not include exon 4 and 5 (~9 copies per cell). The co-localized signals from all three exon-sets and all four sets were exclusively located in the nuclear domain with much lower abundance (**Supplementary Fig. 10**). This is consistent with active *CALU* gene transcription bursts that contain pre-mRNAs in various states of post-transcriptional processing, and thus expected to contain common exons, alternative exons, and introns. In short, by applying RNA FISH, we could reveal the identity and distribution of single mRNA molecules of *CALU* as well as independently confirm the mutual exclusivity of exon 4 and 5 in MCF-7 cells. Together, the results of Sanger sequencing and RNA FISH experiments are in concordance with full-length RNA sequencing and support the coupling events identified.

### Poly(A) signal usage for coupled polyadenylation sites

Most alternative PASs in MCF-7 cells were found in tandem (in the same terminal exon, generating a longer or shorter 3’-UTR). From 5,498 genes with multiple PASs, we identified 10,927 tandem PASs in the same exon across 3,983 genes (72%). From these, 3,465 loci (87%) included PASs that were significantly coupled with alternative TSSs or alternative exons. The high number of tandem 3’ UTRs, in both coupled and uncoupled PAS-exon pairs, partly disagrees with previously reported general shortening of 3’ UTRs in MCF-7 cell line^21^. Nevertheless, many coupling events between alternative PASs and inclusion or exclusion of alternative exons are due to the use of exonic and intronic PASs (8,171 non-tandem PASs), leading to the formation of new 3’ UTRs.

To assess whether certain poly(A) signals are preferentially associated with alternative transcription and splicing, we searched for canonical (AATAAA and ATTAAA) and eleven known non-canonical poly(A) signals in the 35bp sequences upstream of the identified PASs. Canonical poly(A) signals could be found in the 35bp sequences upstream of 54% of all PASs (**Fig. 3A**). The proportion of PASs that could be associated with canonical poly(A) signals was unchanged (55.7%) for those that were coupled with TSSs or alternative exons. However, PASs that were linked with TSSs showed a significantly lower proportion of canonical poly(A) signals (38.6%). Although this decrease was accompanied by a slight increase in known non-canonical poly(A) signals, it was mainly due to the use of alternative PASs for which no known poly(A) signal could be found (**Fig. 3A,B**). This suggests that a novel poly(A) signal and alternative mechanisms may be involved in transcription-coupled polyadenylation in MCF-7 cells. Thus, we screened for enriched motifs in the 35bp sequences upstream of PASs that were not associated with known poly(A) signals. Based on a de novo motif enrichment analysis, we identified the enrichment of AKCCTGG for PASs with unknown poly(A) signal (**Table 1**). This motif was also significantly enriched in PASs that were coupled with alternative TSSs or splicing. Interestingly, the AKCCTGG motif could be associated with the binding site of muscleblind-like (MBNL) protein family, known to play a dual role in the regulation of splicing and polyadenylation^22^,^23^. Each MBNL isoform can bind to slightly different motifs^23^ and a few motifs have been previously associated with MBNL proteins^23-25^. Although all three MBNL proteins are expressed in MCF-7 cells, the enrichment of de novo identified AKCCTGG and the recently reported CWGCMWKS motifs (mainly recognized by MBNL3 protein^23^) were more prominent. Notably, previously identified binding motifs for MBNL1 (CTSCYB^24^ and RSCWTGSK^23^) and MBNL2 (TGCYTSYY^23^) were also enriched in sequences upstream of the PASs without a known poly(A) signal (**Table 1**). However, these motifs were not found to be preferentially associated with PASs that were coupled with alternative TSSs or alternative exons. Together, these results suggest an important role of MBNL proteins in mediating alternative splicing and alternative polyadenylation.

### Identification of binding motifs for RNA-binding proteins potentially involved in coupling

We examined the potential involvement of RNA-binding proteins (RBPs) in the coordination of alternative transcription and mRNA processing events by enrichment analysis of their binding motifs in coupled versus non-coupled exons. We screened three genomic domains relative to donor and acceptor splice-sites of coupled exons for enriched sequence motifs (**Fig. 1E**; also see Methods): the 35bp intronic sequences upstream of the acceptor site (R1), the 32bp exonic sequences downstream of the acceptor sites and upstream of the donor sites (R2), and the 40bp intronic sequences downstream of the donor sites (R3).

For coupled nonterminal exons, the sequences from the R1 domain (upstream of the acceptor) were enriched for motifs (**Table 2**) that can be recognized by the splicing modulators RBM24^26^ and SAMD4A^27^ proteins. In addition, the R2 sequences were enriched for binding sites of RBM4B^28^, NOVA2^29^,^30^, and RBM28^31^ proteins, known to play a role in regulating alternative splicing. In fact, many RBM proteins have been associated with pre- and post-mRNA splicing events. R3 regions (downstream of the donor splice-sites) were also enriched for motifs associated with alternative splicing modulators: FUS^32-34^, SRSF2^35^, RBM5^36-38^, PCBPI1 and PCBPI2^39^,^40^ (**Table 2**). Together, we observed a clear indication that RBPs involved in regulation of alternative splicing and mRNA stability are likely to play a role in preferential selection of alternatively spliced exons.

### Conservation of interdependencies across human tissues

We investigated whether the interdependent transcription, splicing and polyadenylation events identified in MCF-7 cancer cells could also be found in the full-length transcriptomes of three primary human tissues: brain, heart and liver. As the sequencing depth in the primary tissues was lower than that of MCF-7, we could only examine the relatively abundant genes. In human brain, of 5,381 genes that could be assessed for coupling between transcription and mRNA processing, 30.7% were found to have at least one coupling event (**Supplementary Fig. 11A**). In total, we identified 7% of 789,054 possible combinations to be significantly interdependent. Similar patterns could be found in heart and liver, having 25% and 26% of genes exhibiting at least one coupling event, respectively (**Supplementary Fig. 11A**). Pairwise comparison of coupling rates between transcriptomes revealed that the proportion of genes exhibiting coupling in multiple samples show a modest range of 5 – 13%, whereas for genes with multiple full-length transcripts, a significantly larger proportion (ranging from 24 – 40%) exhibit at least one coupling event (**Supplementary Fig. 11B**). Overall, MCF-7 transcriptome showed the largest overlap with other datasets suggesting that by achieving a deeper coverage many more interdependencies may be found that are conserved between tissues.

Next, we assessed the conservation of individual coupling events in different samples. From the total number of feature pairs that were found to be interdependent in at least one sample, by far the majority were found to be specific to a given tissue or MCF-7 cells since only 6 – 14% were found to be coupled in two examined datasets (**Supplementary Fig. 11C**). Interestingly, feature pairs that were found to be interdependent in two samples were generally (~77%) mutually inclusive or exclusive in both tissues (**Supplementary Fig. 11C**). This observation suggests that although most coupling events are tissue- or condition-specific, there seems to be a set of interdependencies that are conserved across multiple human tissues.

### Characterization of the proteome of MCF-7 cells in light of full-length transcripts

The MCF-7 transcriptome consists of 11,350 multi-exon genes and 7,364 polyadenylated single-exon transcripts, from which we could identify open-reading frames (ORFs) for 10,385 and 3,591, respectively, based on full-length mRNA sequences associated with each gene locus (**Supplementary Fig. 12**). While almost every one of the 7,814 multi-transcript genes were found to have coding capacity (97.4%), we could identify ORF for a smaller proportion of 3,536 single-transcript genes (78.4%). In addition, 48.8% of single-exon transcripts were found to have a predicted ORF.

To assess the translational characteristics of MCF-7 cells especially in relation to alternatively spliced isoforms and identified interdependencies between transcription and mRNA processing, we analyzed a publicly available coverage, bottom-up mass-spectrometry (MS) dataset^41^. For MS searching, a customized proteomics search database was constructed, consisting of protein sequences derived from Gencode v19 (95,309 entries), ORFs predicted from MCF-7 PacBio sequences (47,325 entries), and a database of frequently observed contaminant proteins (115 entries). The inclusion of the Gencode database allows for identification of any peptides derived from transcripts that were missed by the PacBio data. The MS searching was done using the Morpheus algorithm^42^, wherein all theoretical peptides resulting from an in silico tryptic digestion of protein entries (from Gencode, PacBio, or contaminants database) were matched against the raw mass spectra to identify peptides. We detected 38,628 unique peptides, passing the FDR of 1%, that could be unambiguously associated with the Gencode (version 19) and/or PacBio-based predicted protein-coding sequences in MCF-7 cells. In 2,872 cases, the identified peptide was only present in the Gencode database, whereas in 358 cases, the peptide was only found in the PacBio database. In addition, we found 2,150 peptides associated with 481 single-exon transcripts.

Identified peptides range from 7 to 56 amino acids (aa) with an average of over 15aa in size (**Fig. 4A**). We observed a strong correlation (r = 0.96; p < 2.2e-16) for the number of peptides per gene based on full-length PacBio or Gencode transcripts (**Fig. 4B**), suggesting that full-length RNA sequencing data can capture a comparable repertoire of protein coding sequences in MCF-7 cells. Still, it is evident that for a few select ultra-long transcripts, such as AHNAK, DYNC1H1 and PLEC, peptides were underrepresented in PacBio database compared to Gencode. This is likely because it is quite challenging to synthesize ultra-long transcripts using current full-length cDNA synthesis protocols. The central domain of AHNAK consists of 128aa repeat that is mostly absent in PacBio multi-exon transcripts and is only reflected as two single-exon transcripts (**Supplementary Fig. 13**). In PacBio data, the DYNC1H1 transcripts are defined in two clusters since no exonic overlap were found between them (**Supplementary Fig. 14**). This leads to a representation of 5’ and 3’ peptide hits in two separate PacBio genes as compared to Gencode. For PLEC transcripts, it is likely that the mRNA molecules arising from these loci are not well represented in PacBio data due to their large size (**Supplementary Fig. 15**). This is also likely the case for the lower abundant genes such as HUWE1 (**Supplementary Fig. 16**).

To scan for potentially novel protein isoforms derived from alternatively spliced transcripts, peptide hits were classified into four groups: single-transcript hits representing peptides associated with gene loci that only have one transcript structure; sub-gene hits representing peptides that are associated with only a subset of transcripts for a given gene; single-gene hits representing peptides that are associated with all transcripts of a given gene; and multi-gene hits that represent peptides associated with transcripts of multiple genes. This classification serves as a measure of specificity for peptide matches (**Supplementary Fig. 17**) as well as evaluating the specificity of transcript annotation in representing alternatively spliced isoforms in MCF-7 cells.

Comparison between Gencode- and PacBio-based classification of peptides revealed that PacBio-based analysis of protein peptides provides a more specific peptide assignment, most likely due to use of a sample-specific set of predicted ORFs from the MCF-7 full-length transcriptome as compared with Gencode annotation (**Fig. 4C,D**; **Supplementary Table 3**). Indeed, Gencode represents protein annotations for the entire human proteome, whereas the MCF-7 derived proteome is specific to breast cancer cells. As shown in the empirical cumulative distribution of relative peptide counts per gene (**Fig. 4C**), there is a clear enrichment of peptides that discriminate between different ORFs derived from the same gene locus in PacBio data compared to Gencode annotation whereas the overall peptide counts per gene remain the same between the two annotations. In fact, 50% of peptides that are classified as single-gene hits (matching all transcripts of the associated gene) were classified as sub-gene hit based on PacBio transcripts due to alternative splicing events that are absent in Gencode version 19 annotation (**Supplementary Table 3**). Likewise, most peptides that were classified as single-transcript hits were either categorized as sub-gene (46%) or single-gene (18%) hits in PacBio.

We identified 358 novel peptide hits that were missed in Gencode (**Fig. 4D**; **Supplementary Fig. 18**) and were mainly represented by sub-gene class of peptides (73%). The quality of these novel peptides was adequate as we observed no difference in the quality of peptide hits between those found only in PacBio (‘ novel’ peptides) versus peptides found in Gencode (‘ known’ peptides), as well as peptides with discordant classification based on Gencode or PacBio (**Supplementary Fig. 19**). Essentially, the distribution of peptide MS search scores (i.e., Q value or Morpheus score) was undistinguishable between each group. However, we found different gene assignments for a set of peptides mainly due to single amino acid substitutions (SAS) in MCF-7 compared to the reference sequence (**Supplementary Fig. 20**). Presence of SAS events without knowledge of their existence through sample-specific protein database can either lead to mismatch or failure to detect peptides using general annotations such as Gencode^43^. This is more prominent for sample-specific differential expression of paralogous genes (**Supplementary Fig. 21,22**). These observations are partly reflected by 41% multi-gene peptides that can be specifically assigned to a single gene using PacBio data.

Although most peptide hits were found within a single exon boundary, 30% were associated with exon-exon junctions covering up to 5 consecutive exons (**Fig. 4E**). From 49,263 peptides derived from parts of the transcripts that span exon-exon junctions, 10,364 peptides were associated with exons that were found to be mutually inclusive as we rarely (< 2%) observed peptides that matched mutually exclusive exons (**Fig. 4E**). As shown for ITGB4 gene (**Fig. 4F**), differential splicing pattern in MCF-7 transcriptome can significantly influence the characterization of matching peptides as well as the impact of coupling in coding protein.

## DISCUSSION

Short-read RNA sequencing has become central in assessing the global RNA expression patterns. However, as a result of the complexity of human transcriptome and limited size of the sequenced RNA fragment, this approach disappoints in precise reconstruction and reliable expression estimation of transcript variants^6^,^7^,^44^. In contrast, single-molecule long-read sequencing provides a unique opportunity to reveal the true complexity of the transcriptome^9^, 10,45,46 as it can determine the full structure of individual transcripts by full-length sequencing.

Here, we have analyzed the deepest and longest transcriptome data so far to better understand the extent of interdependencies between transcription and mRNA processing. Notably, full-length mRNA sequencing and de novo identification of high-quality sequence of transcript variants uncovered an unprecedented amount of potentially novel transcripts in MCF-7 cells and three human tissues. Our findings not only unravel a higher level of alternative transcription, splicing and polyadenylation in MCF-7 transcriptome than previously appreciated, but also provide valuable information on the preferential selection and interdependency between these processes.

We showed that transcription initiation, splicing and 3’ end formation are tightly coupled in over 60% of genes with multiple transcripts and such interdependencies can be found across the entire length of the mRNA molecule. Notably, we report an unforeseen and unprecedented number of genes that undergo a vigorous preferential selection during transcription and mRNA processing as the choice of transcription initiation subsequently influences both alternative splicing of exons and the usage of alternative poly(A) site. Ample evidence points at the critical role for RNA Pol II in the coordination between transcription and mRNA processing (reviewed in 5, 47-49). It has been shown that RNA Pol II initiation, pausing, and elongation rate can influence alternative splicing and polyadenylation of transcripts^50-53^. Moreover, the C-terminal domain of RNA Pol II likely acts as a scaffold for regulatory factors that are involved in splicing and polyadenylation (reviewed in 49). Concordantly, we found an enrichment of coupling events in larger genes that seem to undergo a more extensive regulation during mRNA synthesis. However, the exact mechanisms by which the coordination is achieved remain largely unclear.

From previous studies, it became clear that polyadenylation couples with splicing machinery to influence the removal or inclusion of the last intron^16^,^54^,^55^. We now show that (i) the interdependencies between splicing and polyadenylation are not necessarily restricted to the final introns, (ii) that they can also involve introns that are far from the poly(A) site and (iii) that the coupling between splicing and alternative polyadenylation is not restricted to tandem 3’ UTRs. The exact mechanisms by which these coupling events are achieved fall beyond the scope of this study. Previously, it has been shown that spliceosome components are also part of the human pre-mRNA 3’-end processing complex^56^. Moreover, it is likely that there are RNA-binding proteins with a dual role in alternative splicing and polyadenylation to coordinate mRNA processing events. hnRNP H^18^, CstF64^55^, MBNL1 and ELAV1 (HuR)^22^, ^57-59^ are a few examples of such proteins. Importantly, sequences upstream and downstream of splice-sites (R domains) of coupled exons were enriched for motifs that can be recognized by RNA-binding proteins with known role in regulating alternative splicing. In addition, we found MBNL binding motifs enriched in the sequences upstream of polyadenylation sites coupled with alternatively spliced exons. Interestingly, these regions lacked canonical or non-canonical poly(A) signals. This suggests that MBNL proteins mark alternative poly(A) sites and play a dual and possibly coordinating role in alternative splicing of exons and polyadenylation. This is in line with previous studies in MBNL1-deficient cells where both splicing and polyadenylation were shown to be disrupted^22^, ^23^.

Based on the reported sequence preference of MBNL proteins^23^, MBNL3 is the most likely candidate of the MBNL family responsible for the coordination between alternative splicing and polyadenylation of transcripts in MCF-7 cells. However, it is not clear to what extent these findings can be extrapolated to other cell lines and cell types. In MCF-7 cells, although we generally do not find very long transcripts, we do not see any evidence for a shift to more proximal poly(A) sites that were previously reported^21^,^60^ as many genes have tandem poly(A) sites. Nevertheless, we observe many cases in which the use of alternative poly(A) site is utilized through different 3’ UTRs and not tandem poly(A) sites in the same ’ UTR region. In such cases, the absence of binding sites for regulatory proteins and miRNAs can enhance the tumorigenic activity of MCF-7 cells by allowing transcripts to escape from inhibition^21^. It is not clear whether MBNL-mediated polyadenylation, coupled with transcription initiation and splicing, is achieved through direct recruitment of RNA processing machinery or via alteration of secondary structure and formation of RNA molecules that, in turn, affect the choice for poly(A) site usage. Our analysis also identified a few more candidates with dual roles in mRNA processing, notably multiple RBM proteins SAMD4A, NOVA2, FUS and SRSF2, which warrant further investigations by performing additional functional assays.

In MCF-7 cells, the multitude of protein isoforms arising from alternative transcription and mRNA processing is not fully reflected in Gencode protein annotation as sample-specific set of predicted open-reading frames seem to provide a better specificity in discriminating peptides based on differences in open-reading frames derived from the same gene locus. Furthermore, as shown in this study, the presence of sample-specific single amino acid substitutions can lead to loss or mismatch of peptides when using a generic reference peptide database. Thus, sample-specific set of potentially coding sequences serves as a valuable resource to capture the true complexity of the proteome and to study the global functional divergence between protein isoforms. However, it is important to note that the comprehensiveness of such database vastly depends on the sequencing depth and library preparation strategy and, therefore, it is currently indispensable that such analyses need to be performed using the combination of Gencode and sample-specific protein sequences.

This study demonstrates that our understanding of transcript structures and coordinating mechanisms that regulate transcription and mRNA processing is far from complete, even in well-characterized human cell lines such as MCF-7. Single-molecule full-length RNA sequencing of other human tissues also provide an additional evidence for the true complexity of the human transcriptome. Moreover, although it has been shown that single-nucleotide variants can alter the inclusion of exons in transcripts^9^, it is of interest to identify variants that can affect allele-specific coupling between transcription and mRNA processing. Together, these can offer a better understanding of the mechanisms that control transcription and mRNA processing. As alternative splicing is a key mechanism in functional divergence of human genes, access to full-length sequence of potential protein isoforms allows us to better understand biological function through examining interactions and cross-tissue dynamics of protein isoforms^61^. In turn, this unique set of protein isoform interactions serves as a global view of protein functional repertoire and thereby provide valuable insights into underlying mechanisms of diverging physiological, developmental or pathological conditions.

## METHODS

### RNA sample preparation, library preparation, and sequencing

The methodologies and experimental settings for RNA preparation, cDNA synthesis, library preparation, and sequencing are described at:*https://github.com/PacificBiosciences/DevNet/wiki/IsoSeq-Human-MCF7-Transcriptome*. We downloaded the 2015 dataset, which is an updated version of the original 2013 release. In addition, we used publicly available data from three human tissues (brain, heart and liver) for comparative analysis. These datasets are described at: *http://www.pacb.com/blog/data-release-whole-human-transcriptome/*.

### Annotation of transcripts using isoform-level clustering algorithm (ICE)

The identification, polishing, and annotation of transcripts were previously carried out using the ICE algorithm and made public by Pacific Biosciences^62^.To find transcript clusters, ICE performs a pairwise alignment and reiterative assignment of full-length reads to clusters based on likelihood. This process is followed by consensus calling and further polishing of the sequence to reduce the redundancy and increase the overall accuracy of sequences for identified transcript variants. For further information on the methodology and experimental settingsvisit:*https://github.com/PacificBiosciences/cDNA_primer/wiki*.

### Comparison to the GENCODE annotation

We used GENCODE annotated transcripts (version 19) as reference to compare with the identified transcripts in the human MCF-7 transcriptome data. The comparison was carried out using cuffcompare from the Cufflinks suite^63^.

### Comparison to standard RNA-Seq datasets

We used five publicly available RNA-Seq datasets (SRR1035698, SRR1107833, SRR1107834, SRR1107835 and SRR1313067; generated on Illumina HiSeq2000 or Illumina HiSeq2500 platforms) to evaluate the reliability of gene expression quantification based on full-length mRNA sequencing data used in this study. As accurate transcription reconstruction is not feasible for short-read RNA-Seq data, the comparison is made at the gene level using GENCODE annotation (version 19). Median gene coverage (fragment counts adjusted for gene length) was used as a measure for gene expression quantifications using the GENTRAP pipeline (http://biopet-docs.readthedocs.io/en/latest/pipelines/gentrap/).

### Definition of transcription start site, polyadenylation site, and donor and acceptor splice sites

In this study, by processing the GFF file that contains the annotation of all identified transcripts and exon/intron boundaries (defined by the genomic position and strand on the hg19 reference sequence), a list of all transcription and mRNA processing events is produced. Transcription start sites (TSSs) are defined as the first genomic position of each transcript structure. Polyadenylation sites (PASs) are defined as the last genomic position of each transcript. The most upstream and downstream genomic positions of exons were used to define donor and acceptor splice-sites, respectively. However, for the first exon only the donor site is described as the first position is defined as transcription start site. Likewise, the last exon does not contain a donor splice site as the position is defined as polyadenylation site. If multiple transcripts share the same feature, then only one copy is kept in the unique set of features at each locus. Furthermore, the union of all unique exons is defined as the available sequence at each locus. This is also illustrated in Fig. 1B. Terminal positions of transcripts were curated based on 10bp window to remove stochastic noise and minimize the number of false TSS and PAS for each locus.

### Alignment and quantification of supporting reads for each transcript

The number of reads aligned to each transcript was used as the supporting evidence for each transcript structure. To identify the number of supporting reads, the polished sequences of all unique transcripts were used as a reference for the unique alignment of raw reads using BLASR^64^. Other parameters were set default and according to the PacBio guidelines.

### Statistical analysis

After defining unique features (transcription start sites, exons, and polyadenylation sites) and identifying the number of supporting reads for transcripts at each locus, all possible pairwise comparisons between features were made. To do this, the sum of all reads that support the presence of the two selected features in all observed transcripts is reported in a two-by-two contingency table. The table describes the number of times two features are observed in the same transcript or exclusively, as well as the sum of reads that are mapped to transcripts that do not support the presence of either features (**Fig. 1C**). A significant linkage between two features is assessed using the Fisher’ s exact test. The mutual inclusivity or exclusivity of coupled features are defined using their log-transformed odds-ratio. All p-values are adjusted using Bonferroni multiple testing correction.

Coupling network is constructed based on detected interdependencies between pairs of features for each gene. Nodes represent features and mutual inclusivity or exclusivity is represented by black or red edges, respectively, in the network. Mutual inclusivity or exclusivity sub-networks are constructed after removing all the other edges. No further filtration is performed on gene coupling networks. All statistical analyses were performed in R and Python.

### Sanger sequencing validation

The PCR for Sanger sequence validation was performed using the 2x Phusion High-Fidelity PCR master mix with HF buffer (NEB). Briefly, the pcr ran for 30 cycles with 1 min elongation at 72C. The PCR products were purified using Ampure XP beads following the guidelines of the manufacturer. The sizing of the amplicons was checked using Agilent’ s Labonachip system. The Sanger sequencing of the products was performed by the LGTC and the sequences were analyzed using Sequence Scanner Sofware 2 (Applied Biosystems, CA USA).

### Single molecule RNA fluorescence in situ hybridization

Single-molecule RNA FISH relies on the combined fluorescence from 25-48 singly fluorophore labeled oligonucleotides bound to the same RNA. By using the fluorescence from a guide probe set in one dye, the fluorescence from one or more exon-specific probe sets with < 25 oligonucleotides, and each labeled with a separate dye, can be accurately registered as belong to the same RNA. The optimal number of oligonucleotides per specific set must be experimentally determined.

#### Probe sets

Four probe sets were designed at www.biosearchtech.com/stellarisdesigner to detect: 1) The common exons of the human CALU (Calumenin; NCBI Gene ID: 813; 7q32.1) mRNAs (CALU_E), 2) the alternatively spliced exon 4 (CALU_E4), 3) the alternatively spliced exon 5 (CALU_E5), and the common first intron (CALU_I1). The probe set target sequences were as follows: CALU_E: NM_001199671.1 nts 1-141, 957-1188, 1383-1610, 1811-2875, CALU_E4: NM_001199671.1, nts 1177-1394, CALU_E5:NM_001199672.1, nts 1177-1394, and CALU_I1: NC_000007.14, nts 128739433-128747432. CALU_E is an inclusive probe set designed to detect the following variants: NM_001219.4, NM_001130674.2, NM_001199671.1, NM_001199672.1, NM_001199673.1, and NR_074086.1. Both CALU_E and CALU_I1 are full sets with > 32 oligonucleotides, whereas the sets targeting the short exons 4 and 5 have 9 oligonucleotides each. The four sets were synthesized at LGC Biosearch Technologies as custom Stellaris^®^ probe sets with unique fluorophores: CALU_E: Quasar^®^ 670, CALU_E4: Quasar 570, CALU_E5: Cal Fluor^®^ Red 610, and CALU_I1: FAM. The CALU_E4 and CALU_E5 probes were further purified by reverse phase HPLC, to ensure full labeling.

#### Reagents and smRNA FISH

Human breast adenocarcinoma MCF-7 cells (ATCC-HTB-22) were obtained from ATCC (Manassas, VA) and cultured as recommended by the provider. The hypotriploid karyotype is available at the provider’ s web site, and shows three chromosomal loci for 7q32.1. 2-(4-Amidinophenyl)-6-indolecarbamidine dihydrochloride, 4’ 6-Diamidino-2-phenylindole dihydrochloride (DAPI), molecular biology grade ethanol, acetic acid, and methanol were from Sigma Aldrich (St. Louis, MO). Vectashield was from Vector Laboratories (Burlingame, CA). Stellaris RNA FISH hybridization and wash buffers were from LGC Biosearch Technologies. Stellaris RNA FISH was performed as previously described for methanol/acetic acid-fixed cultured cells^65^,^66^.

#### Image acquisition and analysis

DAPI-stained nuclei, fluorescein (FAM), Quasar 570 (Q570), CalFluor 610 (CF610) and Quasar 670 (Q670) dyes were imaged through a 60X 1.4NA oil-immersion lens on a Nikon TI widefield microscope using the appropriate filters: 49000-ET-DAPI, 49011-ET-FITC, SP102v1, SP103v2, 49022-ET-Cy5.5 respectively. The exposure and sequence of channels to acquire were determined based on the brightness and photostability of the dye with which each probe set was labeled. The sequence of exposure was Q670, followed by FAM, and then either or both Q570, CF610. Each Z-slice was exposed for 1 s, except for Q670 which required 2 s exposures. For each field of view, a range spanning the vertical dimension of the cell (typically 10 um) is defined and for each channel, a series of images were acquired through this span at 0.3 μm increments by using Nikon Elements’ Advanced Research software.

Each Z-series was collapsed and rendered as a single, max-intensity projected image. Translational registration to align images shifted relative to another was accomplished by ImageJ macros after identification of a region containing overlapping signals in each channel. Peak positions of these signals were determined relative to each other to inform the shift of each channel. Next, spots and their centroid positions were identified in each channel using the ImageJ Find Maxima utility. These positions were then compared against one another and co-localized spots were grouped if within 3 pixels (330 nm). Based on these groupings, spots were categorized into separate transcript variants and displayed on the image for review. Finally, cell borders were defined and spots associated with distinct cells for per-cell and per-transcript variant copy number determination. RNA FISH features were counted in at least ten cells.

### Sequence motif analysis relative to polyadenylation sites

For each detected locus, we reported the last nucleotide as polyadenylation site. Each genomic location was converted into a BED format. Strand specific genomic sequences located up to 35 nucleotides upstream each unique polyadenylation site were extracted, in a FASTA format, using UCSC Table Browser (GRCh37/hg19). FASTA files were parsed using a custom bash script to count the number of sequences containing specific 6-mer motifs: one of the two canonical polyadenylation signals AATAAA and ATTAAA, or one of the eleven non-canonical polyadenylation signals (AAGAAA, AATACA, AATAGA, AATATA, AATGAA, ACTAAA, AGTAAA, CATAAA, GATAAA, TATAAA, TTTAAA). Subsequently, the same 6-mer motifs were counted for each unique PAS significantly coupled to TSSs or exons and for each unique PAS that did not show a significant coupling.

For PASs that could not be attributed to known poly(A) signals, we ran DREME67 (v. 4.11.4) to identify enriched motifs. Randomly shuffled set of sequences was generated from the original sequences of the examined PASs and used as a background set. In addition, the sequences of known recognition motifs for MBNL proteins^23-25^ were counted for each set using a custom script. Subsequently, the enrichment of each motif was assessed by Fisher’ s exact test.

### Tandem 3’ UTR analysis

This analysis was performed to identify loci that contain tandem 3’ UTRs (loci that contain more multiple PASs located in the same last exon). Custom scripts were used to identify loci that contain at least two PASs that share the same coordinates of the last exon start. The number of loci with tandem 3’ UTRs was calculated for those in which PAS was significantly coupled to alternative exons and for those that did not show any significant interdependencies between alternative exons and the PAS usage.

### Sequence motif analysis relative to acceptor and donor sites

For each detected locus, we reported the first and last nucleotide of each exon as acceptor splice site and donor splice site, respectively. Each unique genomic position was converted into a BED format and the strand specific sequences of 2 nucleotides length were extracted using UCSC Table Browser (GRCh37/hg19) for both acceptor and donor splice sites. A custom bash script was used to count the number of dinucleotide sequences containing ‘ GT’ and/or ‘ AG’

### RNA binding motif analysis

We used MEME suite tools to identify enriched sequence motifs present in exons significantly coupled with TSSs, PASs or other alternative exons. For each unique exon, three regions were considered: R1 (containing up to 35 nucleotides upstream the acceptor splice site), R2 (containing 32 nucleotides downstream the acceptor splice site and 32 nucleotides upstream the donor splice site),and R3 (containing up to 40 nucleotides downstream the donor splice site). R1, R2 and R3 regions were obtained by extracting strand specific FASTA sequences using UCSC Table Browser (GRCh37/hg19).

We locally ran DREME67 (v. 4.11.4) for each region separately, and performed a motif search using a negative background (R1, R2 and R3 regions from exons that were not significantly coupled). In each case, a maximum of 10 motifs with E-values < 0.05 was reported. The remaining parameters were kept as default. We then compared each motif found by DREME against the human RNA-binding motifs database CISBP-RNA using TOMTOM Motif Comparison tool68. We ran the analysis by setting the Pearson correlation coefficient as comparison function and considered only matches with a minimum false discovery rate (q-values) < 0.05.

### Open reading frame prediction and proteomics data analysis

ORF prediction was done on the PacBio MCF-7 sequences using ANGEL (*http://www.github.com/Magdoll/ANGEL*). Prediction was done on both the PacBio consensus reads and a genome-corrected version of the transcript, and whichever produced the longer ORF was chosen to represent the transcript CDS. The predicted MCF-7 ORF sequences were concatenated with Gencode version 19 and protein sequences representing common mass spectrometry (MS) contaminants, creating a customized FASTA file (i.e., proteomics search database). The Morpheus software (version 131) was employed for MS searching of the custom database against the MCF-7 Thermo Raw files obtained from Geiger et al.^41^ study. Unknown precursor charge state range was set to +2 to +4. Absolute and relative MS/MS intensity thresholds were disabled. Maximum number of MS/MS peaks were set to 400. Assign charge state was set to true. De-isotoping was disabled. The protease specificity was set to trypsin with no proline rule enabled. Up to 1 missed cleavage was allowed and N-terminal methionine truncations was variable. Fixed modifications used were carbamidomethylation of cysteines. Variable modification used was oxidation of methionines. Precursor mass tolerance used was 2.1 Daltons (monoisotopic) and product mass tolerance was 0.025 Daltons (monoisotopic). Modified forms of the same peptide were collapsed and treated as one peptide identification for calculation of false discovery rate (FDR). An FDR of 1% was used to filter for final peptide identifications. All identified peptides were categorized as: single-transcript if the peptide matches to only one gene with one transcript; sub-gene if the peptide matches to a subset of transcripts of only one gene; single-gene if the peptide matches to all transcripts of only one gene; and multi-gene if the peptide matches to multiple transcripts from multiple genes.

### Data availability

Detailed description of the methodologies along with all open-source Python scripts and generated results are made publicly available at: *<u>https://git.lumc.nl/s.y.anvar/mRNA-Coupling/wikis/home</u>*.

## ACKNOWLEDGMENTS

We would like to thank dr. Hagen Tilgner and dr. Stefan White for constructive discussions on the manuscript, dr. Henk Buermans for technical assistance, and Pacific Biosciences for making the full-length mRNA sequencing data available for this study.

## AUTHOR CONTRIBUTIONS

SYA, SWT and PAC’ tH designed the study. SYA, WGA, EdK, MV performed the computational and statistical analyses. ET carried out the identification of unique transcripts, alignment to the genome and open-reading frame predictions. GS performed the proteomics analysis. SYA, ET and GS performed the integration of transcriptomic and proteomic findings. RHY and HEJ carried out the smRNA-FISH experiments. YA performed the Sanger sequencing validation. JTdD and SWT contributed to the experimental design and the interpretation of the findings. SYA and PAC’ tH oversaw the project. SYA wrote the manuscript that was subsequently read and approved by all co-authors.

